# Utilizing Temporal Measurements from UAVs to Assess Root Lodging in Maize and its Impact on Productivity

**DOI:** 10.1101/2020.05.21.108746

**Authors:** Sara B. Tirado, Candice N. Hirsch, Nathan M. Springer

## Abstract

Both stalk and root lodging can cause significant yield losses in maize; however, maize plants are often able to recover from root lodging. There is potential among breeding programs for developing lines that are more tolerant and can more rapidly recover from root lodging. We assessed the incidence of root lodging utilizing end-of-season lodging scores collected among the Genomes 2 Fields (G2F) initiative trials and found a large yet variable incidence of lodging across states, years, and genotypes. Lodging in this dataset was scored manually at the end of the season, and little is known about the drivers of lodging and lodging recovery. We therefore developed an approach for utilizing temporal plant height measurements collected from unmanned aerial vehicles to capture in-season lodging and recovery in a yield trial consisting of 24 maize hybrids planted in replicate under two dates and three planting densities in St Paul, MN in the summers of 2018 and 2019. We found that growth rates during vegetative development as well as the developmental timing of plants when exposed to a storm are predictive of the amount of lodging maize plots will experience. We also found that utilizing temporal height measurements can help in not just estimating lodging and early vegetative growth rates, but that utilizing these estimates can also aid in assessing end of season yield.

## INTRODUCTION

Lodging has strong negative impacts on maize yield due to reductions in the photosynthetic capacity of plants, the inability to mechanically harvest lodged plants that did not recover, and the rotting of ears due to being in contact with the ground. Lodging also limits the application of ground-based vehicle navigation through fields by causing the blockage of rows. Lodging occurs primarily through two avenues, one taking place at the plant stalk and the other at the root. Annual yield losses attributable to stalk lodging are estimated to be 5-20%; however, losses for root lodging are not as well documented (Flint-Garcia et al., 2003; Zuber & Kang, 1978). The specific type as well as the severity of the lodging event that occurs can be dependent on many factors including the developmental stage of the crop when experiencing the storm, the severity and patterning of the wind, and a variety of morphological characteristics of the genotype being grown including the stalk strength, root depth and root branching (Ma et al., 2014; Robertson et al., 2014; Sanguineti et al., 1998).

Lodging events that occur at the stalk, referred to as snapping, are most prominent during maize plants’ vegetative growth stages when internodes are growing thereby rapidly weakening the cell walls leading to stalk breakage if exposed to strong winds or at the grain-filling stage (Ching et al., 2010). Stalk lodging hinders plant development if it occurs in vegetative stages and also ear development if it occurs in reproductive stages and plants are unable to recover. Root lodging, however, most often occurs in response to early-season storm events before maize plants have fully developed brace roots (Ennos et al., 1993). At this point, plants have more plasticity and have the ability to recover from the lodging event through a mechanism referred to as ‘goose-necking’ (Zhang et al., 2011). Even partial recovery to the upright orientation through goose-necking can help minimize yield losses (Dhugga, 2007). This leads to opportunities in breeding pertaining to both root lodging resistance as well as root lodging recovery.

Genotypic resistance to root lodging has been addressed by breeding improved hybrids over the last decades with consistently higher lodging tolerance (Duvick, 2005). However, a large portion of a plant’s lodging reaction is dependent on environmental factors as well as interactions between the genotype and the environment making lodging resistance a complex trait to study. Root lodging in particular occurs very irregularly across environments as it is influenced by a multitude of factors such as soil quality and fertility, wind patterns and pest and disease pressures making it challenging to maintain consistent progress in selection (Flint-Garcia et al., 2003; Stamp & Kiel, 1992). To aid in studying root lodging responses, several traits have been utilized to indirectly measure lodging resistance including the horizontal pushing resistance, root pulling resistance, root volume by water replacement, root clump size and weight, and more recently root failure moment (Ennos et al., 1993; Fouéré et al., 1995; Kamara et al., 2003; J. Liu et al., 2011; S. Liu et al., 2012; Thompson, 1968). These however are destructive and labor-intensive measurements. A different approach that has been taken to evaluate lodging resistance has been to control wind conditions by using a mobile wind machine (Barreiro et al., 2008; Wen et al., 2019). Most studies, however, rely on irregular weather events and subsequently measuring root lodging by visually inspecting and manually annotating the number of plants leaning more than 30 or 45 degrees from vertical. Because this is time-consuming, lodging measurements are typically scored at one timepoint at the end of the season prior to harvest rather than following extreme weather events. These measurements do not capture plants that lodged but were able to fully recover their upright orientation without producing a significant bend in the lower portion of the stalk. Although this is useful in determining genotypes that suffered a higher degree of lodging throughout the season, it gives little insight into how genotypes respond in terms of lodging throughout development and how the timing of the lodging event impacts plant recovery and end-season productivity.

The incidence and severity of lodging responses is exacerbated by the increased incidence of extreme weather events that accompany climate change, and developing tools and knowledge around the drivers of lodging and recovery will allow us to mitigate risk and breed superior germplasm. Assessing the degree of in-season lodging and the rates of recovery require temporal data collection and unmanned aerial vehicles (UAVs) offer a potentially fast and cost-effective way of evaluating fields remotely in an automated fashion. The goals of this study were therefore to develop an approach for measuring lodging responses in a fast and effective manner utilizing UAVs, to use this pipeline to evaluate management and developmental factors that contribute to the severity of lodging responses observed for a set of hybrid lines grown under six different environment-by-management conditions in MN, and to test the impacts of lodging severity as well as response and rate of recovery on yield at the end of the season.

## RESULTS

### Root Lodging Incidence Across the US

The Genomes to Fields (G2F) initiative has generated a publicly available large-scale, multi-year, and multi-state dataset of maize phenotypes. Using this dataset we sought to evaluate the frequency of lodging for varying environments and genotypes ((McFarland et al., 2020)). Environmental and phenotypic data for approximately 74,000 field plots involving more than 2,500 maize hybrid varieties evaluated across twenty states in North America between 2014-2018 has been released (McFarland et al., 2020). Among other traits, the incidence of root lodging, measured as the number of plants per plot where the stalk was 45 degrees or more from vertical as maturity, as well as plot grain yield extrapolated to bushels per acre was collected as part of this experiment. Twenty total states collected phenotypic data throughout the five years; however, there were multiple field sites that did not collect the necessary data for assessing lodging and only 11 out of the 20 states had lodging and stand count data for all five years.

Of the 73,686 G2F plots grown from 2014 to 2018 with lodging data, 10.9% of these suffered from root lodging (based on >5% of all plants within the plot being scored as lodged) revealing a large incidence of lodging across the US in recent years, with extremely variable quantities across years (Figure 1A). An analysis of variance (ANOVA) showed a significant genotypic and environmental contribution to the percentage of plants in a plot that experienced root lodging (Table S1). To better assess the variability of lodging responses, we classified plots that lodged into two categories (partial and extreme lodging) based on the percentage of plants per plot that lodged (Figure S1, see methods). There is substantial variation in the percentage of plots that experienced root lodging both across states in individual years and across years in individual states (Figure 1B). Field sites in some locations, such as Ohio and Texas, experienced very small degrees of root lodging across all years. In contrast, field sites in other states, such as Iowa, experienced large degrees of root lodging throughout all years. There were six states that experienced root lodging in at least one plot across all five years. Other sites, such as Nebraska and Illinois, were very variable and experienced a high degree of lodging some years but no lodging in other years. This variation in lodging responses is likely due to a combination of differences in climatic conditions across states and years as well as in differences in the management practices implemented in different locations, such as the planting date and density.

**Figure 1.**
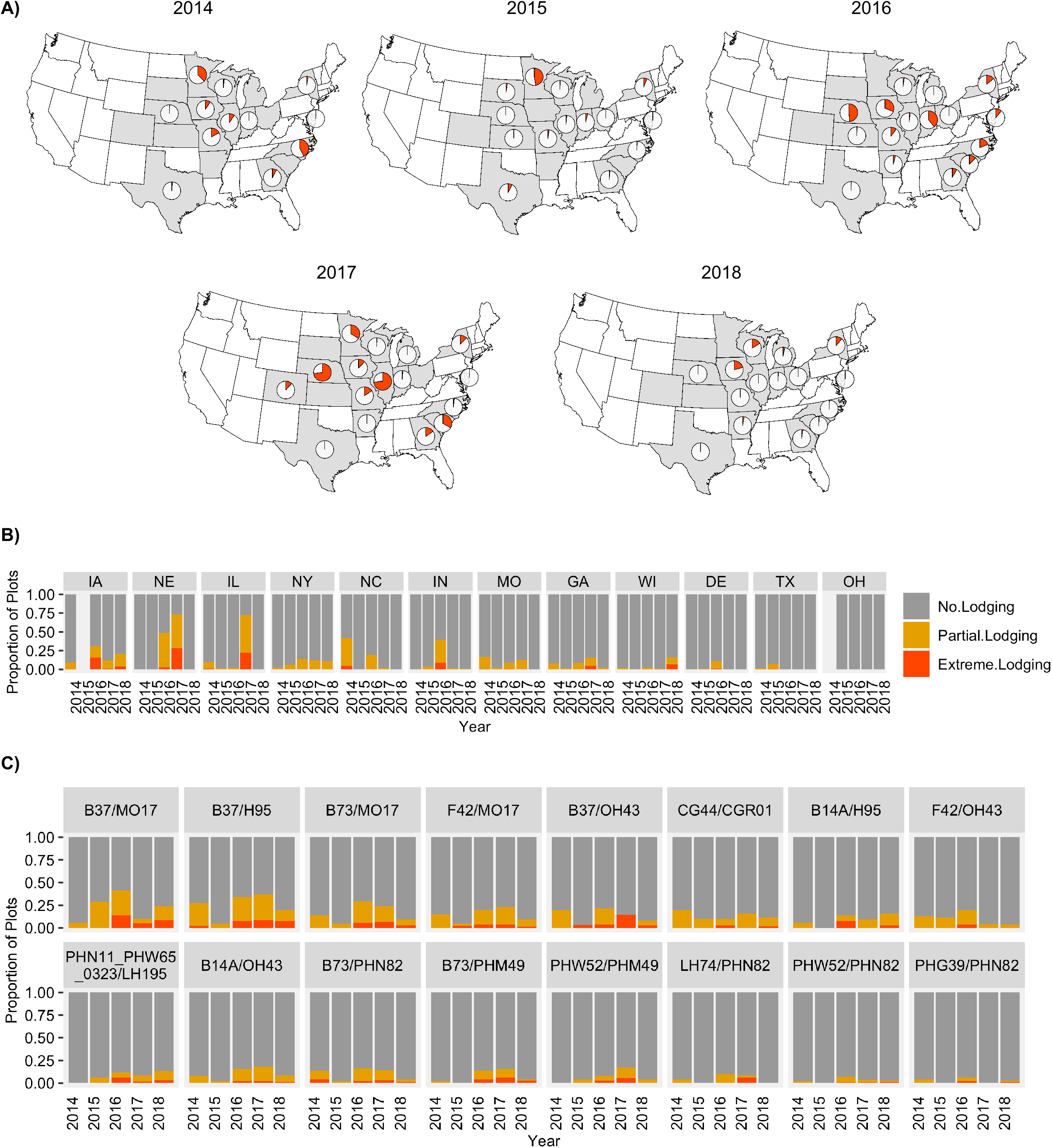
**A)** Percent of plots in G2F locations across years with lodging data where more than 5% of plots lodged. **B)** Percent of plots per state across all years where less than 5% of plants lodged (grey), between 5 and 50% of plants lodged (yellow) and more than 50% of plants lodged (red). **C)** Percent of plots across all locations and years for the 16 genotypes with lodging data in at least 6 locations all five years where less than 5% of plants lodged (grey), between 5 and 50% of plants lodged (yellow) and more than 50% of plants lodged (red).

Past studies have found variation in lodging resistance based on genotypic background (Landi et al., 2007; Nelimor et al., 2020). The G2F dataset also provides an opportunity to assess genetic variation for lodging. There was large variability in the amount of genotypes that lodged in at least one environment across the five years ranging from 337 out of 1206 genotypes in 2015 to 714 out of 1161 genotypes in 2018. We focused our analysis on a small subset of genotypes that were planted at many (>6) locations with lodging data available each season (Figure 1C). Half of the genotypes experienced lodging in at least one location all five years; however, both the total percentage of plots and the severity of the lodging responses observed varied by genotype (Figure 1C). A subset of the genotypes, such as B37xMo17 or B37xH95, exhibit consistently higher levels of lodging across each year. Other genotypes, such as PHG39xPHN82 or PHW52xPHN82, consistently exhibit very low levels of lodging. The end of season lodging observations within the G2F dataset highlights the prevalence of lodging. However, from this data there is a lack of information about the specific events that trigger lodging or data on the ability of genotypes to recover from these events.

### Measuring Lodging Responses Using UAVs

To further our understanding of the factors that contribute to variability in lodging from different events we utilized data from two growing seasons in which naturally occurring weather events provided an opportunity to study variability in lodging as well as the rate and extent of recovery. These experiments consisted of a set of 12 hybrid genotypes grown in replicate 4-row plots with two planting dates (mid May and late May), three planting densities (60k/ha, 90k/ha and 120k/ha) and two years (2018 and 2019) in St Paul, MN (Figure S1A). The 2018 dataset consisted of 13 timepoints of UAV data collection, whereas the 2019 dataset consisted of 23 timepoints (Figure S1B). In both years, a storm event with strong winds occurred during the vegetative phase of development (July 1st in 2018 and July 15th in 2019) and caused extensive root lodging. Planting densities and sowing dates have been shown to influence end-season root lodging scores across maize genotypes (Li et al., 2015; Sher et al., 2017). The design of these experiments with multiple planting dates and densities enable us to study the impact of developmental stage, planting density, and genotypic variation on the extent of initial lodging, post-lodging recovery, and impacts on end of season traits including height and yield.

The ability to track lodging and lodging recovery responses requires measurements taken at high temporal resolutions, which is often not feasible with manual observation. UAVs on the other hand provide an efficient means to monitor plant responses to extreme wind events since they can collect data in a fast and efficient manner at high temporal resolutions (Y. Shi et al., 2016). UAV plant height (PH-UAV) measurements were collected at many timepoints throughout development for both of these experiments including the day prior to the wind event and the day following the wind event (Figure S2). The UAV-derived digital elevation models (DEMs) of plots on the date prior to the lodging event and the DEMs of the day after were compared and a reduction in height across certain plots in the field was observed (Figure 2A-B). This prompted us to develop a method to use PH-UAV measurements to quantify lodging. To develop a lodging severity score from the PH-UAV measurements, the percent change in average plot height extracted from the DEM from the closest timepoint before the storm to the closest timepoint after the storm was calculated (see methods). The severity of the lodging response (no lodging, moderate lodging, or severe lodging) was then determined for each plot based on this percent change in height (Figure S1; Figure 2B; see methods). The ability to apply this method to determine lodging is influenced by the timepoints of data collected before and after the lodging event. Due to the lower temporal resolution of UAV data collection in the 2018 season, the timepoint closest to the lodging event was 3 days prior, whereas data was collected the day before the lodging event in the 2019 season (Figure S2). For both years, flights were conducted the day following the lodging event. Because this approach involves calculating lodging based on the percent change in plot height before to after the storm event, we highlight the importance of having high temporal resolution in UAV measurements to be able to capture plot height at a date close before a storm occurs.

**Figure 2.**
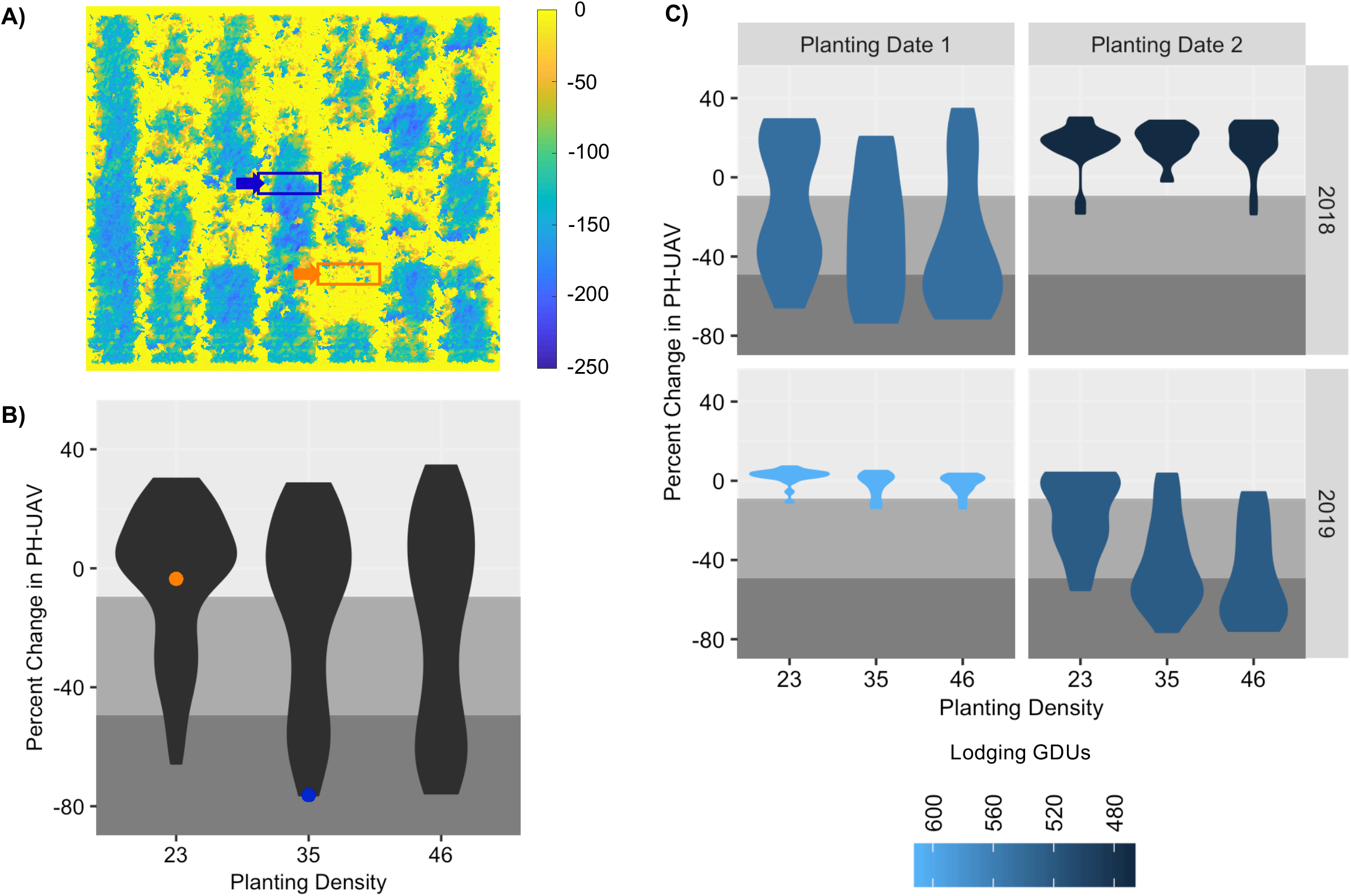
**A)** Height difference (in cm) for the 2019 late planting date treatment at 7/16/2019 compared to 7/15/2019 following the wind storm that occurred the night of 7/15/2019. Boxes represent plots with low (orange) and high (blue) degrees of lodging. **B)** Density plot of lodging percent height change for all plots across all treatments in the 2018 and 2019 seasons. The background shading represents the areas where plots where depicted as having no height change (light grey), moderate height change (medium grey), and severe height change (dark grey). Points represent plots marked in panel A with low (orange) and high (blue) degrees of lodging. **C)** Density plot of lodging percent height change for all plots across individual planting date treatments and years. The color represents the growing degree units (GDUs) of the specific treatment-year combination at the time of the storm event.

### Variation in Lodging Responses

Substantial variability in lodging responses was observed across plots (Figure 2A; Figure 2B). Within environments, there were also major differences in the amount of lodging observed for the two planting dates in each season as well as more subtle differences based on planting density (Figure 2C). The differences in lodging between the two planting dates suggest that plots that were either early in vegetative development or nearing the reproductive stage of development did not experience large amounts of lodging. The early planting had more substantial lodging in 2018 and the later planting had more substantial lodging in 2019; however, these plantings were at similar growing degree units (GDUs) when the lodging event happened (Figure 2C). This suggests that maize hybrids are most susceptible to root lodging within a narrow developmental timeframe, and were less impacted in this case when younger than 469 GDUs or older than 590 GDUs.

The developmental timepoint and growing conditions at the time of the storm impacted not just the amount of lodging experienced by certain plots, but also their rates of recovery from the lodging event. For example, LH82 x LH145 grown at high density had a much slower recovery rate when grown in 2018 in the first plating date treatment compared to when grown in 2019 in the second planting date treatment, even though both experienced similar amounts of lodging (Figure 3). It took six days for the 2018 plot to get back to the original height it was before the storm event, compared to only 3 days in 2019. The slower recovery rate in the lodged early planting 2018 plot impacted its terminal height achieved and it ended up having a much shorter terminal PH compared to the lodging event that occurred later in development in the 2019 plot (Figure 3). The difference between the terminal PH of the lodged plot compared to the non or semi-lodged counterpart grown in the opposite planting date for each year was a lot larger in 2018 compared to 2019 (Figure 3). It is also worth noting that the 2019 plot grown in the first planting date treatment suffered a small amount of lodging and it was able to recover its original height in just one day. This slight lodging and rapid recovery is something we would have missed in 2018 due to having lower temporal resolution of UAV flights.

**Figure 3.**
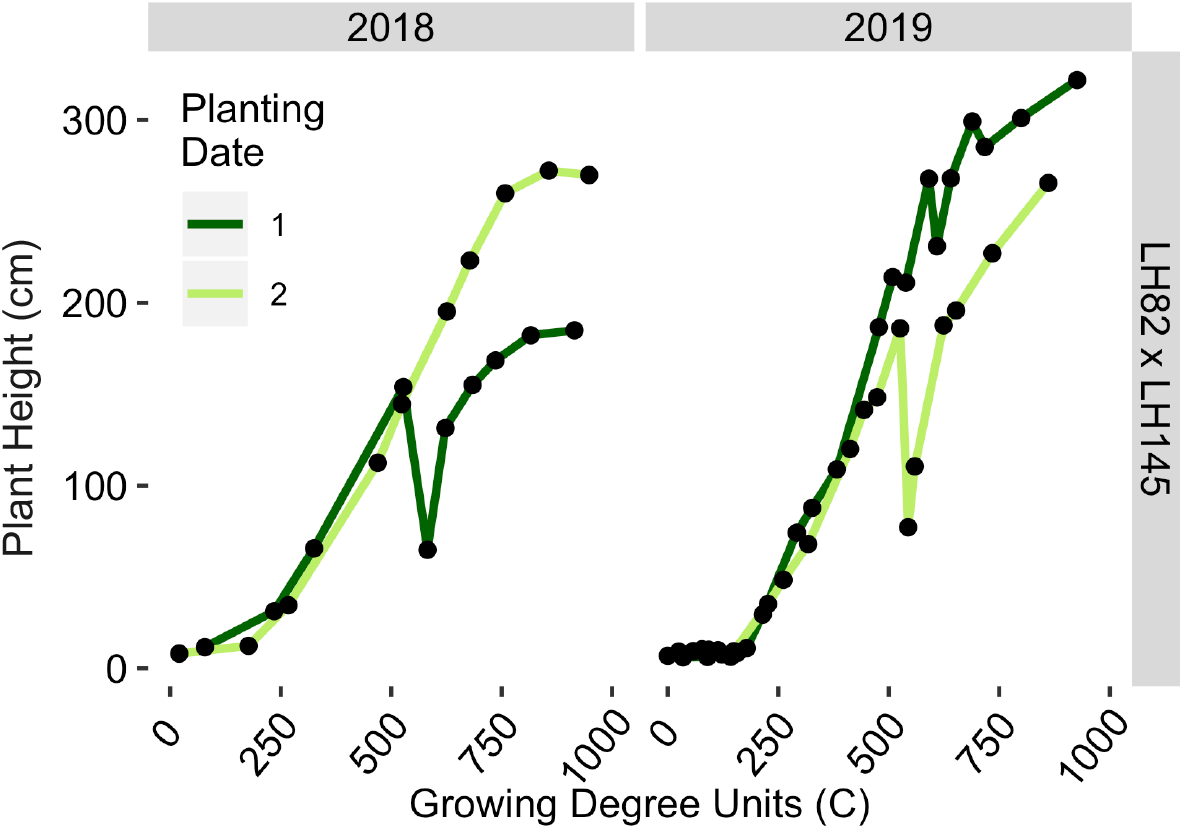
Growth through time for genotype LH82 x LH145 planted at high density across both planting date treatments in the 2018 and 2019 seasons. Points indicate UAV height collection events for each season.

Our analyses suggest substantial variation in lodging that can be attributed to a variety of factors. The different genotypes, planting dates and densities all appeared to influence the lodging responses observed across both seasons (Figure 4A; Figure S3). An ANOVA was performed to assess the contributions of different factors to explain the change in height observed by UAV measurements (Table 1, Table S2). Multiple factors including the genotype, the planting density, and the plot height prior to the lodging event significantly contributed to the variation in lodging responses observed across both years in the planting date that suffered high degrees of lodging (Table 1). The higher temporal resolution in 2019 allowed us to calculate the slope, or growth rate, across early and mid-season timepoints preceding the lodging event (Figure 4B; see methods). These factors were included in the 2019 ANOVA and reveal that mid-season growth rates also significantly contributed to the amount of variation observed for lodging severity (Table 1). Our findings suggest that growth rates together with the developmental time point a plot is in at the time of a storm event will largely dictate the severity of the initial lodging event.

**Table 1.** ANOVA results for lodging percent height change and yield for plots in the 2018 first planting date treatment and in the 2019 second planting date treatment.

| 2018 P1 | Lodging Height Change |  |  |  | Yield |  |  |  |
| --- | --- | --- | --- | --- | --- | --- | --- | --- |
| Variable | Df | Sum Sq | Pr(>F) | Signif | Df | Sum Sq | Pr(>F) | Signif |
| Genotype | 11 | 19983 | 0.0049 | ** | 11 | 0.12861 | 0.012887 | * |
| Density | 1 | 4199 | 0.0120 | * | 1 | 0.05869 | 7.94E-04 | *** |
| Replicate | 1 | 1552 | 0.1184 | N.S. | 1 | 0.0038 | 3.55E-01 | N.S. |
| EarlyG | NA | NA | NA |  | NA | NA | NA |  |
| MidG | NA | NA | NA |  | NA | NA | NA |  |
| PreLodgingPH | 1 | 15147 | 1.01E-05 | *** | 1 | 0.00061 | 0.70994 | N.S. |
| Lodging | NA | NA | NA |  | 1 | 0.00631 | 0.235375 | N.S. |
| Recov3 | NA | NA | NA |  | 1 | 0.00145 | 0.566106 | N.S. |
| Recov6 | NA | NA | NA |  | 1 | 0.00061 | 0.710188 | N.S. |
| TerminalPH | NA | NA | NA |  | 1 | 0.0034 | 0.381359 | N.S. |
| Genotype:Density | 11 | 7570 | 0.3662 | N.S. | 11 | 0.09571 | 0.058598 | . |
| Residuals | 45 | 27563 |  |  | 34 | 0.14706 |  |  |
| 2019 P2 | Lodging Height Change |  |  |  | Yield |  |  |  |
| Variable | Df | Sum Sq | Pr(>F) | Signif | Df | Sum Sq | Pr(>F) | Signif |
| Genotype | 11 | 22871 | 5.10E-15 | *** | 11 | 0.029278 | 5.02E-03 | ** |
| Density | 1 | 12022 | 3.34E-15 | *** | 1 | 0.002051 | 0.1277 | N.S. |
| Replicate | 1 | 7 | 0.7686 | N.S. | 1 | 0.000845 | 0.32327 | N.S. |
| EarlyG | 1 | 1 | 0.9012 | N.S. | 1 | 0.000004 | 0.94378 | N.S. |
| MidG | 1 | 5512 | 3.86E-10 | *** | 1 | 0.004187 | 0.03246 | * |
| PreLodgingPH | 1 | 547 | 0.0147 | * | 1 | 0.000313 | 0.54572 | N.S. |
| Lodging | NA | NA | NA |  | 1 | 0.001206 | 0.23943 | N.S. |
| Recov3 | NA | NA | NA |  | 1 | 0.003828 | 4.03E-02 | N.S. |
| Recov6 | NA | NA | NA |  | 1 | 0.000062 | 0.78738 | N.S. |
| TerminalPH | NA | NA | NA |  | 1 | 0.01909 | 3.65E-05 | *** |
| Genotype:Density | 11 | 617 | 0.7660 | N.S. | 11 | 0.008398 | 0.54267 | N.S. |
| Residuals | 43 | 3646 |  |  | 33 | 0.027716 |  |  |
| '.' significant at 0.1, '*' significant at p=0.05; '**' significant at p=0.01; '***' significant at p=0.001; 'N.S.', not significant. |  |  |  |  |  |  |  |  |

**Figure 4.**
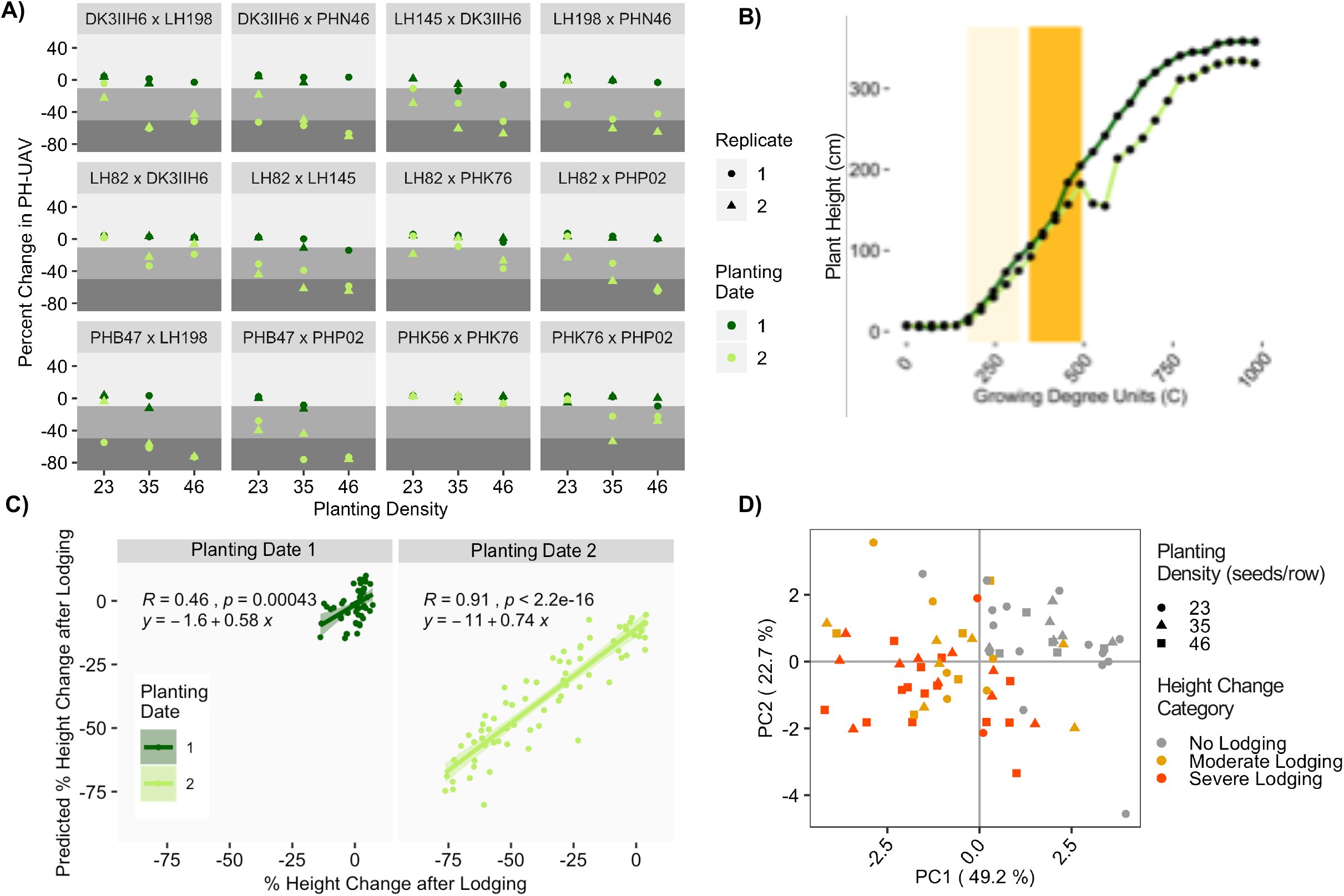
**A)** Lodging percent height change for replicate plots of each genotype in the 2019 season. The background shading represents the areas where plots were depicted as having no lodging (light grey), moderate lodging (medium grey), and severe lodging (dark grey). **B)** Height through time for a single replicate of a single genotype (LH145 x DK3IIH6) planted at high density in the first and second planting date treatments. The background shading represents the time ranges selected to calculate the early (light yellow) and mid-season (dark yellow) growth rates. **C)** Lodging percent height change predictions. The prediction model contained planting density, PH prior to the lodging event, early-season growth rate, and mid-season growth rate as the predictor variables. **D)** PC1 and PC2 scores from the PCA of the slope between each timepoint within the early and mid-season growth periods shown in panel B of all predicted height values derived from the loess fitted model for all 2019 plots. Points are colored by lodging height change category and shaped by planting density.

### Predictors of Lodging Responses

Genotypic resistance to root lodging has mostly been addressed through indirect selection of morphological traits found to be highly correlated with lodging resistance, and this has driven efforts to identify QTL linked to these traits (Bruce et al., 2001; Landi et al., 2007; J. Shi et al., 2019). The ability to more efficiently assess in-season root lodging opens new possibilities for identifying new phenotypes linked to lodging resistance that can be selected for and improved.

Moreover, identifying correlated phenotypes that can be more easily assessed utilizing UAVs can improve the resolution of genotypic selection efforts by allowing breeders to sample larger panels of genotypes across more locations (Araus & Cairns, 2014; Y. Shi et al., 2016). We sought to determine what factors contribute to the lodging responses observed and whether there are variables that could be strong predictors for lodging. Using the 2019 dataset due to its higher temporal resolution, we ran a stepwise model selection procedure on a linear regression model that incorporated multiple variables (see methods) to predict the lodging percent height change. The selected model contained the planting density, the PH prior to the lodging event, the early-season growth rate, and the mid-season growth rate as the predictor variables. This model, derived from half of the dataset, predicted the lodging height change on the other half of the dataset with very high accuracy (adj R-squared of 0.46 for the first planting date and 0.91 in the second planting date; Figure 4C). The lower accuracy in the first planting date was likely due to a small degree of PH change across most plots due to limited amounts of lodging. These results indicate the importance that rates of growth across early vegetative stages as well as height values achieved by plants at the time when they are exposed to storm events have in determining the degree of lodging.

Rates of plant growth can be driven by how genotypes respond to environmental conditions. In this case, the environment encompasses a mixture of management treatments including planting date and planting density together with temperature, precipitation and other climatic factors. Studies have shown that planting density affects plant growth by influencing the ratio of root to shoot growth, the rate and quantity of root biomass accumulation, and the rate of internode elongation thereby affecting both plant and ear height as well as lodging (Hébert et al., 2001; Jiang et al., 2013; Tetio-Kagho & Gardner, 1988; Xue et al., 2016). Because the key drivers we found to be strong indicators of lodging were related to PH and rate of change in PH prior to the lodging event, we hypothesized that these traits were influenced by the response of each genotype to the density treatments they were exposed to. To test this, a lowess model was fit across timepoints for each plot in the 2019 season and the plot height across consistent timepoints throughout development for each planting date was extracted with a temporal resolution of 35 GDUs. The slope between each timepoint within the early and mid-season growth rate periods was calculated and used as input into a principal component analysis. The results show that the first two principal components (PC1 and PC2) were able to separate non-lodged from lodged plots as well as planting density with higher planting densities aligning along the axis of higher lodging severities (Figure 4D). Slopes across the latest timepoints within the mid-season growing period just prior to the storm event had the largest loadings for PC2 and the slope for timepoints within the transitioning phase between early and mid-season growth had the largest loadings for PC1. Plots also clustered by genotype suggesting that the genotype by density component had an impact in lodging severity as well by modulating the PH response and rate of growth of individual plots (Figure S3).

### Lodging Impact on End-season Productivity

Seasonal yield predictions have become very important across agricultural stakeholders such as farmers, commodity traders and government officials for making strategic decisions with regards to management and planning. Yield predictions have been done through many methods including surveys, statistical models and process-based models and studies have been executed to more efficiently predict yield across many crops utilizing a multitude of information sources including climatic data, physiological and phenological data as well as remote sensing data (Basso & Liu, 2019). Most studies that have utilized remote sensing data, however, have relied on single-timepoint remote-sensing measurements and haven’t investigated the use of temporal measurements in yield predictions. We therefore wanted to investigate the contribution of different factors obtained through temporal measurements including early-season lodging and factors that contributed to lodging, such as early-season and mid-season growth rates, in predicting final yield. Root lodging has already been shown to contribute to grain yield in maize lines exposed to heat and drought stress treatments and to be useful in identifying lines resistant to abiotic stresses (Nelimor et al., 2020). Scoring in-season lodging and lodging recovery throughout the season therefore shows potential to improve yield predictions and germplasm selection.

We first looked at the G2F plots with end of season lodging information. A strong, significant impact of root lodging and the degree of lodging severity scored by hand at the end of the season on yield was observed (Figure 5A). Lodging led to a 12.3 bu/acre yield loss when comparing plots that experienced extreme lodging to those that experienced no lodging.

**Figure 5.**
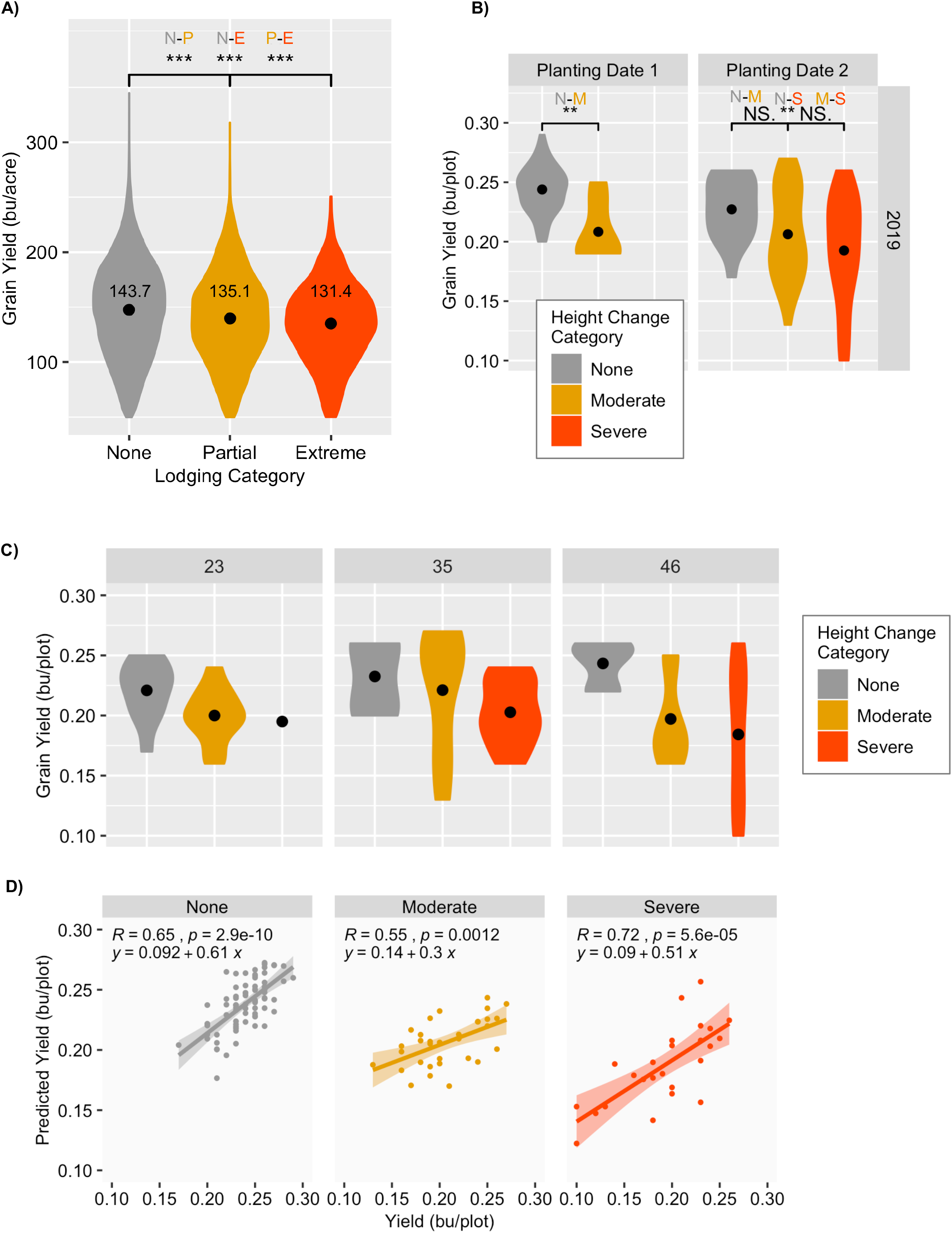
**A)** Density plot of grain yield (in bushels/acre) for all the Genomes 2 Fields Initiative hybrid plots with lodging data grown across all states and years grouped by lodging score category (none, partial and extreme). Significance values based on t-test comparing different groups are plotted (* significant at p=0.05; ** significant at p=0.01; *** significant at p=0.001; N.S., not significant). **B)** Density plot of grain yield (in bushels/plot) for all the 2019 experimental plots grouped by height change category (none, moderate and severe) and by planting date. Significance values based on t-test comparing different the groups are plotted (* significant at p=0.05; ** significant at p=0.01; *** significant at p=0.001; NS., not significant). **C)** Density plot of grain yield (in bushels/plot) for all the 2019 experimental plots in the second planting date treatment grouped by height change category (none, moderate and severe) and by planting density (23, 35, and 46 seeds/row). **D)** Grain yield predictions split by height change category on 1/2th of all 2019 plots using a model generated with the other 1/2th of the dataset compared to the measured yield values. The model contained genotype, early-season growth rate, lodging height change, and terminal PH as the predictor variables.

Similarly, lodging percent height change extracted in-season led to significant yield declines in our experimental plots (Figure 5B). This yield decline was most prominent in high-density planting treatments compared to lower density treatments (Figure 5C).

To better assess which factors were most predictive for yield and to determine the relative impact of lodging on end-season productivity, we again ran a stepwise model selection procedure on a linear regression model that incorporated multiple variables to predict end-season yield for the 2019 dataset (see methods). The variables selected and used to create the yield prediction model were the genotype, the early-season growth rate, the lodging height change, and the terminal PH. This model was able to explain between 55% and 72% of the variation observed for yield across the different lodging severity groups (Figure 5D). This indicates the utility of temporal measurements in yield predictions and the potential of incorporating remote-sensing temporal data together with weather and agronomic data for yield prediction model generation. This would, however, require enough temporal resolution to capture unexpected responses such as lodging and storm recovery events.

## DISCUSSION

Assessing the degree of in-season root lodging and the rate of recovery is important for better understanding root lodging resistance and for finding suitable lines for environments that are prone to lodging. Evaluating root lodging data collected for thousands of genotypes grown throughout many US locations across multiple years as part of the Genome 2 Fields initiative showed that there are a large number of plots that have lodged across the US in recent years. However, the number and severity are extremely variable across years and states. There is also substantial genetic variation for lodging among the genotypes included in this dataset. For this dataset lodging was only evaluated at the end of the season and therefore does not capture the variable ability of genotypes to recover from mid-season lodging events.

To better track lodging responses, we utilized UAV-derived plant height data collected temporally to assess how different genotypes planted at two timepoints (an early and a late planting) and grown under various planting densities (low, medium and high density treatments) across two years respond to storm events. To measure lodging and lodging recovery, we developed a metric based on the percent change in height for each plot in response to the storm. Although our metric would capture both root and stalk lodging responses, our plots primarily suffered from root lodging since the storm events occurred across both years during vegetative development when plants are most susceptible to root lodging. If distinguishing between root and stalk lodging was needed, recovery rates could be used to partition both since plants are able to recover to some extent from root lodging but not stalk lodging.

Root lodging responses within our experiment showed that early vegetative growth rates as well as the precise developmental phase of maize plants when exposed to a storm event will determine their response in terms of lodging severity and their rate of recovery. Our maize hybrids were most susceptible to root lodging within a narrow developmental timeframe and plots that were either 469 GDUs or younger or 590 GDUs or older were less impacted and did not experience high degrees of lodging. There was also a genotype by density component modulating early vegetative growth rates which in turn were predictive of lodging responses.

Lodging had a significant, negative yield impact on both the G2F hybrids as well as our experimental plots. Root lodging can cause yield to decrease by reducing the photosynthetic capacity of plants, limiting ear collection from a combine, as well as causing grain mold on ears that were in contact with wet soil. The genotype together with early season growth rates, lodging responses, and terminal height explained a large portion of the variation observed for plot yield. This shows the ability to use temporal measurements derived from UAVs to capture plant responses to weather events to aid in finding lodging resistant lines and determining field productivity in terms of yield. We also note that the lodging events that were tracked in the experimental sites in Saint Paul as well as many of the G2F sites would preclude the ability to use ground-based phenotyping platforms. In many of these field sites the degree of lodging, at least for some of the plots, was sufficient to block the ability of a ground-based vehicle to move through rows. While UAVs are limited in their ability to monitor traits below the canopy they do provide the ability to survey fields that are no longer navigable by ground.

## METHODS

### Lodging Score Estimation for the G2F Dataset

Genome 2 Fields data from 2014 to 2018 was downloaded from https://datacommons.cyverse.org/browse/iplant/home/shared/commons_repo/curated/GenomesToFields_2014_2017_v1 (McFarland et al., 2020). The percentage of plants per plot within the Genome 2 Fields dataset that suffered from lodging was calculated by dividing the number of plants per plot recorded as experiencing root lodging at the end of the season by the total stand counts. These percentages were then assigned a categorical score based on their value (no height change if <5%, moderate height change if between 5% and 50%, and severe height change if >50%; Figure S4).

Variation in percent of plants per plot that experienced root lodging was partitioned into genotype, state, year, replicate, and residual. All factors were treated as fixed effects with the linear model y_ijkl_ = u + g_i_ + e_j_ + y_k_ + r_k_ + p_ijk_ + (ge)_i*j_ + e_ijkl_ where y_ijk_ is the phenotype value of the i^th^ genotype in the j^th^ state in the k^th^ year of the l^th^ replication; u is the phenotypic mean across states and years; g_i_ is the i^th^ genotype effect; e_j_ is the j^th^ state effect; y_k_ is the k^th^ year effect; r_l_ is the l^th^ replication effect; (ey)_j*k_ is the interaction effect of the j^th^ state by the k^th^ year; and e_ijkl_ is the residual effect. To test the significance of the various effect variables of the linear model on lodging height change, we used the anova function of the R stats package (Team & Others, 2013).

### Experimental Field Design

Two replicates of 12 maize hybrids were planted in 4-row plots at two dates (early and late planting) and three densities (60k, 90k and 120k plants per hectare) following a Randomized Complete Block Design blocked by planting date and replicate nested within planting date in Saint Paul, MN in the summers of 2018 and 2019 (Figure S1A). The hybrids utilized were DK3IIH6 x LH198, DK3IIH6 x PHN46, LH145 x DK3IIH6, LH198 x PHN46, LH82 x LH145, LH82 x PHK76, LH82 x PHP02, PHB47 x LH198, PHB47 x PHP02, LH82 x DK3IIH6, PHK56 x PHK76, and PHK76 x PHP02. Each row was 15ft long center-to-center with 12 ft of plots and 3ft of alleys and 30in spacing between rows. There were a total of 144 four-row plots planted each year. The 12 hybrid genotypes utilized were generated by crossing exPVP lines selected due to their past use in production settings and the availability of seed. This experiment was grown on two acres. Nine 1ft x 1ft ground targets in the form of PCV crosses were distributed around the border and internal alleys of the field for use as ground control points (GCPs) based on previously developed optimization algorithms (Gómez-Candón et al., 2014); Figure S5).

### UAV Data Collection and Processing

UAV data was collected approximately weekly from plant germination to plant physiological maturity. A total of 13 timepoints were collected in 2018 and 23 timepoints in 2019, with 2019 having a much higher temporal resolution during the vegetative growth stages in plant development (Figure S1B). UAV data was processed following the procedure established by (Tirado et al., 2019) to extract DEMs and average plot plant height values for each date of data collection. The scripts and processes utilized to perform the image analyses and trait extraction are available at https://github.com/SBTirado/UAV_PH.git.

### UAV Lodging Percent Height Change Estimation

Lodging percent height change (L_n_HC) for each plot each year was measured utilizing the equation

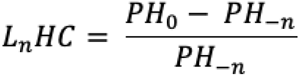

where PH_0_ is the mean plot plant height extracted from a given plot the day following the storm event and PH_-n_ is the mean plot plant height extracted from a given plot at the closest timepoint (*n* days) of UAV data collection prior to the storm event. The values of n corresponded to three and one day prior to the storm for 2018 and 2019 respectively.The lodging percent height change categories of each plot were then determined based on the L_n_HC values (no height change if L_n_HC > −10%, moderate height change if −10% < L_n_HC > −50%, and severe height change if L_n_HC < −50%; Figure S4)

### UAV Lodging Recovery Percent Height Change Estimation

Lodging recovery percent height change (R_n_HC) for each plot each year was measured utilizing the equation

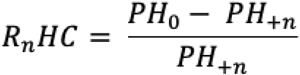

where PH_0_ is the mean plot plant height extracted from a given plot the day following the storm event and PH_+n_ is the mean plot plant height extracted from a given plot at the closest timepoint (*n* days) of the lodging data collection event. The values of n corresponded to three and six for the 2018 season and to one, three, four and six in the 2019 season.

### Weather Data and Growing Degree Days Calculation

Temperature data from an in-field weather station was gathered. Because the weather station each year was installed a few days after planting, daily min and max temperature data from the University of Minnesota St Paul weather station (Station ID 218450) was extracted for these first days after planting (Minnesota Department of Natural Resources, 2020). Growing Degree Units (GDUs) in were then calculated for each date utilizing the equation

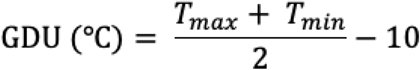

where GDU is the growing degree units accumulated for a single day and T_max_ and T_min_ are the maximum and minimum recorded air temperature values in degrees Celsius for the given date. Temperature values above 30◻ were adjusted to 30◻ and likewise values below 10◻ were adjusted to 10◻ since corn growth rates do not increase or decrease outside the range of these values. A factor of 10l was used as the base temperature and subtracted to the temperature difference to calculate the final daily GDUs. The cumulative sum of GDUs for data collection dates was then extracted as the sum of GDUs for all dates between the planting date and the given date.

### Early-season and Mid-season Growth Rate Extractions

A loess regression model with a span of 0.3 was fit to the average UAV-derived PH of all time points for each plot in the 2019 season as this model was able to accurately capture lodging responses but reduce noise as to create a growth curve. Height for 35 GDUs within the range of 0 to 1000 were then predicted using the fitted model to obtain height across even time intervals for plots across both planting date treatments. Correlation for height across timepoints prior to the lodging event revealed two groups of correlated timepoints and these were selected to calculate the early-season and mid-season growth rates by taking the change in height from the first to the last GDU within each time period and dividing it by the number of GDUs. The early-season growth rate period ranged from 175 to 315 GDUs and the mid-season growth rate period ranged from 350 to 490 GDUs.

### Yield Data Collection

In 2019, grain from the middle two rows of each plot was harvested with a Kinkaid 8XP plot combine equipped with a Harvest Master Single High Capacity GrainGage with Mirus harvest software and total grain weight, test weight, and the percent moisture were collected after all plants had reached physiological maturity. Grain yield per plot was then adjusted to 15.5% grain moisture.

### Statistics

Variation in plot lodging percent height change values was partitioned into genotype, density, replicate, pre-lodging PH, and genotype-by-density interaction, and residual. All factors were treated as fixed effects with the linear model y_ijk_ = u + g_i_ + e_j_ + r_k_ + p_ijk_ + (ge)_i*j_ + e_ijk_ where y_ijk_ is the phenotype value of the i^th^ genotype in the j^th^ planting density of the k^th^ replication; u is the phenotypic mean across planting densities; g_i_ is the i^th^ genotype effect; e_j_ is the j^th^ planting density effect; r_k_ is the k^th^ replication effect; p_ijk_ is the effect of PH prior to the lodging event of the i^th^ genotype in the j^th^ planting density of the k^th^ replication; (ge)_i*j_ is the interaction effect of the i^th^ genotype by the j^th^ planting density; and e_ijk_ is the residual effect. To test the significance of the various effect variables of the linear model on lodging height change, we used the anova function of the R stats package (Team & Others, 2013).

Variation in plot yield values was partitioned into genotype, density, replicate, early and mid-season growth rates, pre-lodging PH, lodging height change, recovery height change three and six days after the lodging event, terminal PH, genotype-by-density interaction, and residual. All factors were treated as fixed effects with the linear model y_ijk_ = u + g_i_ + e_j_ + r_k_ + S^e^_ijk_ + S^m^ + p^1^+ h^3^_ijk_ + h^6^_ijk_ + p^2^_ijk_ + (ge)_i*j_ + e_ijk_ where y_ijk_ is the phenotype value of the i^th^ genotype in the j^th^ planting density of the k^th^ replication; u is the phenotypic mean across planting densities; g_i_ is the i^th^ genotype effect; e_j_ is the j^th^ planting density effect; r_k_ is the k^th^ replication effect; S^e^_ijk_ is the effect of the early-season growth rates of the i^th^ genotype in the j^th^ planting density of the k^th^ replication; S^m^ is the effect of the mid-season growth rates of the i^th^ genotype in the j^th^ planting density of the k^th^ replication; p^1^_ijk_ is the effect of PH prior to the lodging event of the i^th^ genotype in the j^th^ planting density of the k^th^ replication; h^3^_ijk_ is the effect of the recovery height change 3 days after the lodging event of the i^th^ genotype in the j^th^ planting density of the k^th^ replication; h^6^_ijk_ is the effect of the recovery height change 6 days after the lodging event of the i^th^ genotype in the j^th^ planting density of the k^th^ replication; p^1^_ijk_ is the effect of terminal of the i^th^ genotype in the j^th^ planting density of the k^th^ replication; (ge)_i*j_ is the interaction effect of the i^th^ genotype by the j^th^ planting density; and e_ijk_ is the residual effect. To test the significance of the various effect variables of the linear model on yield, we used the anova function of the R stats package (Team & Others, 2013).

### Model Development for Predicting Lodging Severity

A model was built for using various explanatory variables to predict lodging percent height change values for each plot. An optimal model was selected at each time point based on an original model containing the variables genotype, plating density, pre-lodging PH, lodging GDUs, early-season growth rate, and mid-season growth rate using a stepwise model selection procedure with a 10-fold cross validation step. This was implemented using the train function of the caret R package and selecting the model with the lowest RMSE (Kuhn et al., 2016). These predictor variables were chosen based on their biological foundation as well as their level of significance for predicting lodging percent height change values. The selected model contained the planting density, PH prior to the lodging event, early-season growth rate, and mid-season growth rate as the predictor variables.

### Model Development for Predicting End-Season Yield

A model was built for using various explanatory variables to predict yield values for each plot. An optimal model was selected at each time point based on an original model containing the variables genotype, plating density, pre-lodging PH, lodging GDUs, early-season growth rate, mid-season growth rate, lodging percent height change, reovey height change three and six days after the lodging event, and terminal PH using a stepwise model selection procedure with a 10-fold cross validation step. This was implemented using the train function of the caret R package and selecting the model with the lowest RMSE (Kuhn et al., 2016). These predictor variables were chosen based on their biological foundation as well as their level of significance for predicting lodging percent height change values. The selected model contained the genotype, early-season growth rate, lodging percent height change, and terminal PH as the predictor variables.

## ABBREVIATIONS

DEM: Digital Elevation Model
G2F: Genome 2 Fields
PH: Plant Height
UAV: Unmanned Aerial Vehicle

## Data Availability

All data including the UAV-derived plot height values, DEMs and plot boundary files for each date of UAV data collection, cumulative GDUs calculated for each date of data collection, and yield data has been made available at the Digital Repository for U of M.

## ACKNOWLEDGMENTS

We would like to thank Amanda Gilbert and Pete Hermanson for technical support for this experiment. We would also like to thank Anna Deneen, Kjell Sandstrom, Shale Demuth, Danielle Sorensen and Jordan Freeman for helping collect and process the UAV data for this experiment.

## FUNDING SOURCES

This work was supported by the Minnesota Corn Research and Promotion Council. S.B.T. was funded by the University of Minnesota Graduate Opportunity Fellowship, the University of Minnesota APS Metric Funds Fellowship, and a Monsanto/University of Minnesota Multifunctional Agriculture Initiative Graduate Student Fellowship.

## SUPPLEMENTAL FIGURES

**Figure S1**. Categorical scoring system for determining plots that suffered from different lodging severities based on hand measured lodging scores (top) and UAV-derived lodging height change measurements (bottom).

**Figure S2**. **A)** Field experimental layout for 2018 and biological material. **B)** Resolution of UAV imagery collection for the 2018 (orange) and 2019 (blue) seasons with the timepoints following the lodging event of each year represented in red.

**Figure S3**. Lodging severity for replicate plots of each genotype in the 2019 season. The background shading represents the areas where plots where depicted as having no lodging (light grey), moderate lodging (medium grey), and severe lodging (dark grey).

**Figure S4**. PC1 and PC2 scores from PCA of all predicted height values derived from the loess fitted model from the slope between each timepoint within the early and mid-season growth periods colored by genotype.

**Figure S5**. Ground control point placement throughout field border and internal alleys.

## SUPPLEMENTAL TABLES

**Table S1**. ANOVA results for the percentage of plants within a plot with root lodging for all plots in the Genomes 2 Fields dataset from 2014 to 2018 without missing data for plants that experienced root lodging, stand counts and yield measurements.

**Table S2**. ANOVA results for lodging percent height change and yield for plots in the 2018 second planting date treatment and in the 2019 first planting date treatment.

## Notes

### Competing Interest Statement

The authors have declared no competing interest.

