## Supplemental Material for "Utilizing Temporal Measurements from UAVs to Assess Root Lodging in Maize and its Impact on Productivity"

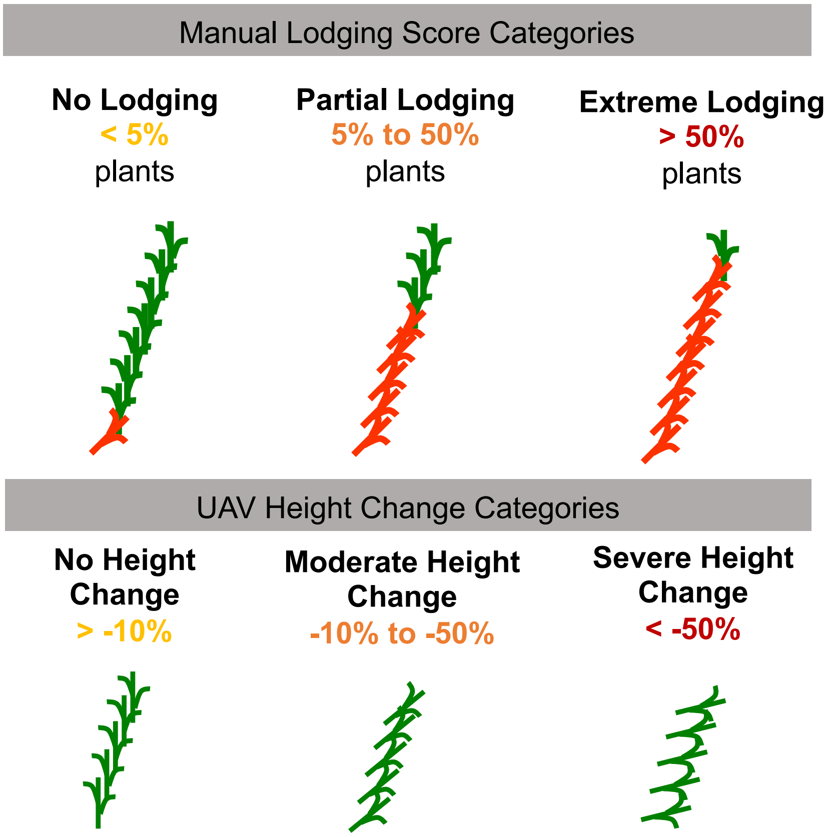


**Figure S1**. Categorical scoring system for determining plots that suffered from different lodging severities based on hand measured lodging scores (top) and UAV-derived lodging height change measurements (bottom).

| | Degrees of Freedom | Sum of Squares | Pr(>F) | Significance |
|---|---|---|---|---|
| Genotype | 2444 | 51.5 | <2e-16 | *** |
| State | 17 | 31.1 | <2e-16 | *** |
| Year | 3 | 6.3 | <2e-16 | *** |
| Replicate | 1 | 0 | 0.387 | N.S. |
| State:Year | 28 | 78.5 | <2e-16 | *** |
| Residuals | 34417 | 452.8 | 1.01E-05 | |

'.' significant at 0.1, ' * ' significant at p=0.05; ' ** ' significant at p=0.01; ' *** ' significant at p=0.001; ' N.S. ', not significant.

**Table S1**. ANOVA results for the percentage of plants within a plot with root lodging for all plots in the Genomes 2 Fields dataset from 2014 to 2018 without missing data for plants that experienced root lodging, stand counts and yield measurements.

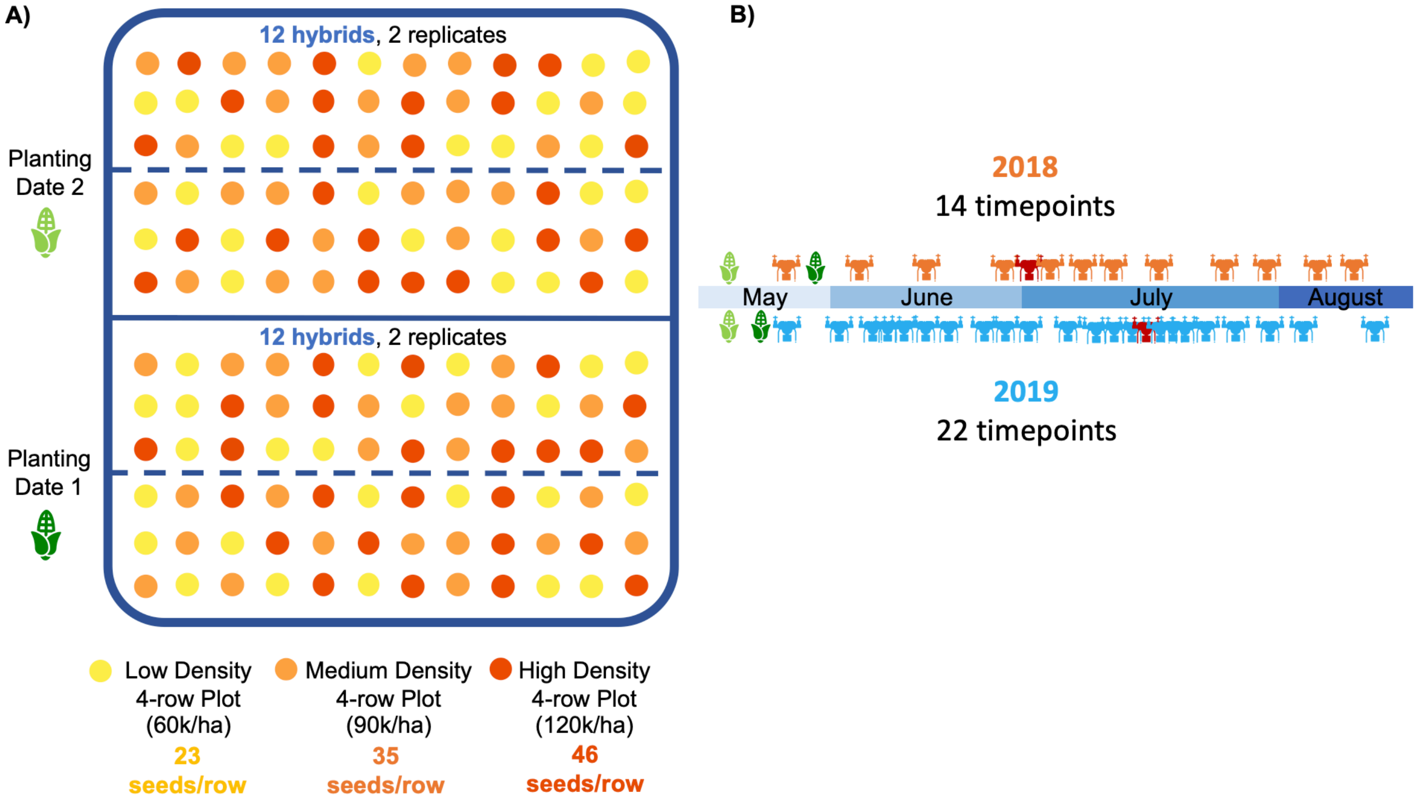


**Figure S2**. **A)** Field experimental layout for 2018 and biological material. **B)** Resolution of UAV imagery collection for the 2018 (orange) and 2019 (blue) seasons with the timepoints following the lodging event of each year represented in red.

| 2018 P2 | Lodging Height Change | | | | Yield | | | |
|---|---|---|---|---|---|---|---|---|
| Variable | DF | Sum Sq | Pr(>F) | | DF | Sum Sq | Pr(>F) | |
| Genotype | 11 | 1294.2 | 0.0792 | . | 11 | 0.08686 | 0.0011 | ** |
| Density | 1 | 0.1 | 0.9638 | | 1 | 0.03726 | 1.43E-04 | *** |
| Replicate | 1 | 852.1 | 0.0007 | *** | 1 | 0.00966 | 3.58E-02 | * |
| EarlyG | NA | NA | NA | | NA | NA | NA | |
| MidG | NA | NA | NA | | NA | NA | NA | |
| PreLodgingPH | 1 | 963 | 0.0004 | *** | 1 | 0.00461 | 0.1401 | |
| Lodging | NA | NA | NA | | 1 | 0.00163 | 0.3744 | |
| Recov3 | NA | NA | NA | | 1 | 0.00451 | 0.1440 | |
| Recov6 | NA | NA | NA | | 1 | 0.00176 | 0.3575 | |
| TerminalPH | NA | NA | NA | | 1 | 0.01034 | 0.0302 | * |
| Genotype:Density | 11 | 1533.3 | 0.0354 | * | 11 | 0.02731 | 0.3057 | |
| Residuals | 44 | 2843.1 | | | 33 | 0.06651 | | |
| **2019 P1** | **Lodging Height Change** | | | | **Yield** | | | |
| **Variable** | **DF** | **Sum Sq** | **Pr(>F)** | | **DF** | **Sum Sq** | **Pr(>F)** | |
| Genotype | 11 | 631.9 | 0.0008 | *** | 11 | 0.0234 | 1.27E-02 | * |
| Density | 1 | 292.7 | 6.97E-05 | *** | 1 | 0.000139 | 0.6712 | |
| Replicate | 1 | 21.7 | 0.2272 | | 1 | 0.000226 | 0.5887 | |
| EarlyG | 1 | 0 | 0.9563 | | 1 | 0.015159 | 0.0001 | *** |
| MidG | 1 | 0.1 | 0.9201 | | 1 | 0.004632 | 0.0194 | * |
| PreLodgingPH | 1 | 5.2 | 0.5502 | | 1 | 0.001023 | 0.2543 | |
| Lodging | NA | NA | NA | | 1 | 0.004043 | 0.0281 | * |
| Recov3 | NA | NA | NA | | 1 | 0.002058 | 0.1099 | |
| Recov6 | NA | NA | NA | | 1 | 0.000435 | 0.4545 | |
| TerminalPH | NA | NA | NA | | 1 | 0.002386 | 0.0862 | . |
| Genotype:Density | 11 | 278.4 | 0.1005 | | 11 | 0.010942 | 0.2658 | |
| Residuals | 35 | 503.6 | | | 29 | 0.021937 | | |

* significant at p=0.05; ** significant at p=0.01; *** significant at p=0.001; N.S., not significant.

**Table S2**. ANOVA results for lodging percent height change and yield for plots in the 2018 second planting date treatment and in the 2019 first planting date treatment.

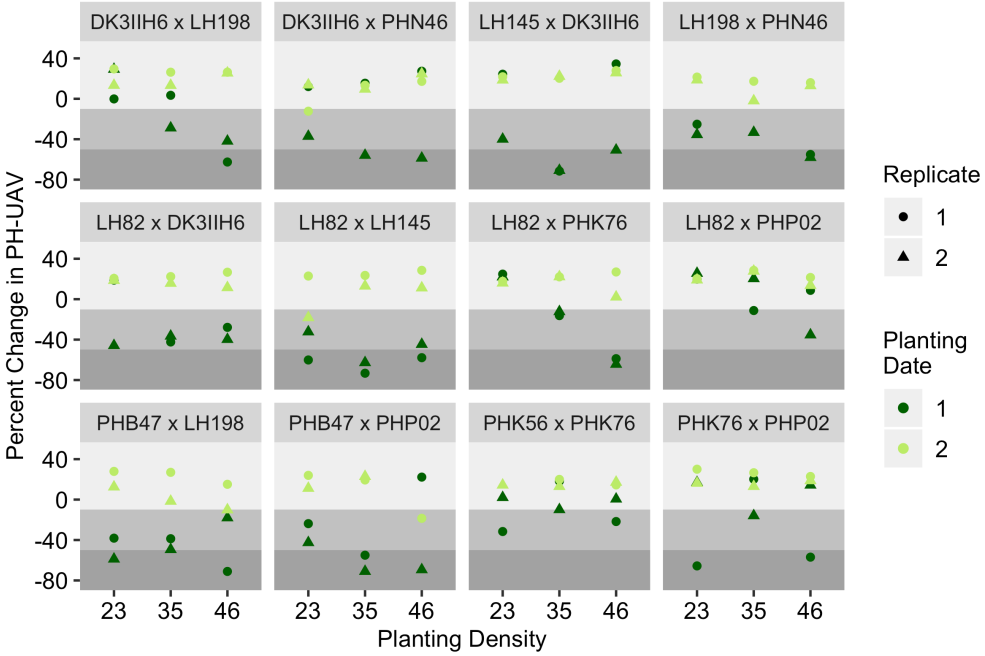


**Figure S3**. Lodging severity for replicate plots of each genotype in the 2019 season. The background shading represents the areas where plots where depicted as having no lodging (light grey), moderate lodging (medium grey), and severe lodging (dark grey).

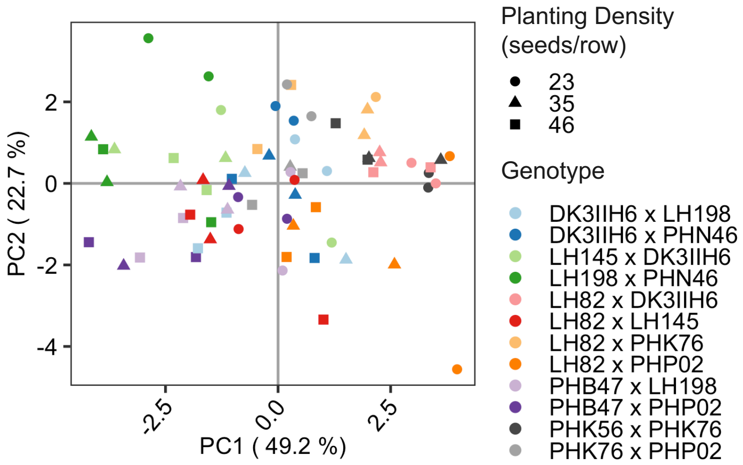


**Figure S4**. PC1 and PC2 scores from PCA of all predicted height values derived from the loess fitted model from the slope between each timepoint within the early and mid-season growth periods colored by genotype.

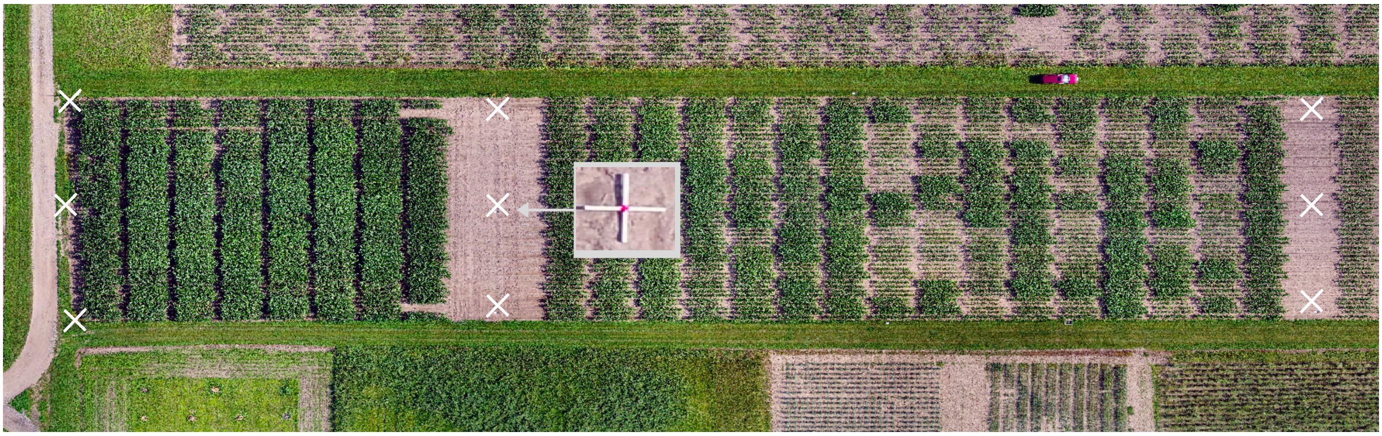

**Figure S5**. Ground control point placement throughout field border and internal alleys.
